# Sex-specific dynamics of activity-dependent regulation of synaptic CaMKII

**DOI:** 10.64898/2026.10.21.760771

**Authors:** Ankit Sharma, Kelsey J. Clements, Natalie D’Ambra, Julia Birnbaum, Elena A. Kuklin, Martha L. Reed, Leslie C. Griffith

## Abstract

Similar behavioral outcomes in males and females can mask substantial differences in the mechanisms underlying neuronal function. However, to what extent sex influences the abundance and activity-dependent changes of proteins that govern neuronal plasticity remains poorly understood. Here we investigated sex-specific regulation of calcium/calmodulin-dependent protein kinase II (CaMKII), an evolutionarily conserved regulator of synaptic stability and plasticity in fruit fly *Drosophila melanogaster*. We find that the steady state levels and activity-driven changes in CaMKII protein differ between female and male at both third instar larval neuromuscular junction (NMJ) and in adult fly mushroom body (MB). These sex-specific differences are downstream of the sex determination pathway, implicating somatic sex identity in the regulation of CaMKII abundance and its activity-dependent dynamics. Despite the differences in CaMKII levels between female and male flies, we observed no impairment of plasticity in either sex. Overall, our findings reveal sex-dependent regulation of both steady-state and activity-dependent levels of a highly conserved neuronal plasticity-associated protein across distinct neuronal contexts and development stages. These results further support the idea that females and males can achieve similar behavioral outcomes through different molecular mechanisms. Growing evidence that sexually dimorphic molecular responses contribute directly to sex differences in vulnerability to stress and disease underscores the importance of identifying these mechanisms and understanding their consequences.

**Significance Statement:** Neuronal sex differences are commonly studied in reproductive circuits or when neuronal structure or function differs overtly between males and females. However, similar physiological and behavioral outcomes may hide distinct underlying molecular mechanisms. Here, we show that both steady-state CaMKII protein levels and activity-dependent changes in CaMKII differ between female and male flies at the larval neuromuscular junction and in adult mushroom body neurons. These differences are downstream of the sex-determination pathway but occur without overt changes in behavioral output. Our findings reveal a hidden layer of sexual dimorphism in the dynamic regulation of synaptic proteins, demonstrating that equivalent function need not reflect equivalent molecular mechanisms. Such differences may contribute to sex-specific vulnerabilities to neurological and psychiatric disorders.

## Introduction

Sexual dimorphism in the nervous system has been most thoroughly studied in the context of sex-specific behaviors underlying reproduction (Meeh et al., 2021; Tong et al., 2026). It is becoming increasingly clear, however, that sex differences occur at every level of neuronal function, including in the molecular pathways that generate behaviors whose overt output does not differ between males and females. Understanding sexual dimorphism at this mechanistic level has importance for human health, as most neurological and psychiatric disorders exhibit clinically relevant sex differences in prevalence, presentation, progression, outcome, or treatment response (Mauvais-Jarvis et al., 2020).

The pathways underlying learning and memory provide a striking example. The dominant mechanistic model of memory formation, rodent hippocampal long-term potentiation (LTP), was developed overwhelmingly in males (Gall et al., 2023) and was initially assumed to apply to females without direct testing. This began to change when investigators found that LTP varied across the estrous cycle in females (Warren et al., 1995). Subsequent work identified distinct pathways underlying LTP in males and females (Jain et al., 2019; Copenhaver and LeGates, 2024; Le et al., 2024) indicating that measuring plasticity alone may not capture sex-dependent differences in neuronal signaling. Rather, the same functional outcome may emerge from different molecular or cellular processes in females and males (Hyer et al., 2018; Gall et al., 2023).

Such differences are not restricted to vertebrates (Salzberg et al., 2023). Although the mechanisms that establish sex-specific circuitry in flies and worms are increasingly well understood, much less is known about sex-dependent differences in intracellular signaling and plasticity within circuits that are not overtly sexually dimorphic. RNA seq studies using *C. elegans* have suggested that there are substantial sex biases in the expression of many genes in neurons shared between females and males (Haque et al., 2025). In *Drosophila* the story is less clear since most of the comparative profiling has been done from neuronal subsets that demonstrate structural differences between males and females (Chen et al., 2026). Nonetheless, the precedent from worms and mammals suggests the baseline state of neurons in females and male flies are likely to be different as a consequence of biases in gene expression.

Sex differences in plasticity may also arise from differences in the spatial and temporal dynamics of shared signaling pathways driven by posttranscriptional regulation. Because the amplitude, duration, and localization of signaling determine downstream cellular responses, differences in pathway dynamics can produce distinct mechanisms of plasticity even when the starting state or the final physiological or behavioral outcome is similar. These types of difference are of particular interest to human health since they can result in differential vulnerabilities in females and males. Where disease-relevant sex-dependent regulation of cellular pathways has been studied in detail, it has often been found to be based on evolutionarily conserved molecular dimorphisms (Pomatto et al., 2017; Buck et al., 2021; Regan et al., 2022).

In this study, we ask whether calcium/calmodulin-dependent protein kinase II (CaMKII), a key and evolutionarily conserved regulator of synaptic plasticity, exhibits sex-specific differences in is abundance and activity-dependent regulation. CaMKII is a highly conserved serine/threonine protein kinase enriched in nervous system of both flies and vertebrates and is a central regulator of neuronal plasticity (Nicoll and Schulman, 2023). Importantly, CaMKII protein levels are themselves strongly regulated by activity in both rodents (Miller et al., 2002) and flies (Nesler et al., 2016; Clements et al., 2026), making dynamics a key aspect of its function. We show that basal and behavior-stimulated CaMKII protein differs between females and males at both the third-instar larval neuromuscular junction (NMJ) and in adult mushroom body (MB) neurons. This difference is specified by the cell-autonomous sex-determination factor *Transformer (tra)*. Despite these striking differences in CaMKII regulation, we detect no corresponding impairment of plasticity in either sex. Together, these findings support the idea that males and females can achieve similar plastic outcomes through differently organized molecular mechanisms and protein dynamics.

## Materials and Methods

### Fly lines and maintenance

All fly stocks were cultured on standard food at room temperature. Experimental flies were kept at 25°C and 70% relative humidity on a 12 h light, 12 h dark period. Fly lines used in this study include: VT030559-GAL4 (206077 VDRC), *C380-Gal4* (BL80580; Presynaptic driver), *MHC-Gal4* (BL38464; Postsynaptic driver), UAS-GCaMP6f, UAS-Flp,(BL4539) FRT::STOP::FRT-GFP-CaMKII (Chen et al. (2022), UAS-Tra RNAi (a generous gift from Prof. Michelle Arbeitman).

### Larval NMJ

#### 5x Spaced depolarization/high potassium stimulation

Third instar wandering larvae were chosen for experiments to ensure they were at the same developmental stage. Sex of the larvae were identified by the presence (male) or absence (female) of visible gonads. Larvae were dissected in cold normal HL3.1 and then checked to ensure that the axons and central nervous system were intact. Larvae were kept unstretched so that muscles had room to contract during stimulation. Larvae were stimulated as described in Ataman et al. 2008; they were incubated in high potassium HL3.1 5 pulses of 2 min, 2 min, 2 min, 4 min, 6 min each, with rest in between each pulse and at the end in normal HL3.1 The molarity and components of normal and high K+ HL3.1 were the following: Normal HL3.1: 70 NaCl, 5 KCl, 0.1 CaCl_2_, 10 MgCl_2_, 10 NaHCO_3_, 5 trehalose, 107 sucrose, 5 HEPES-NaOH, pH 7.2; and high K+ HL3.1: 40 NaCl, 85 KCl, 1 CaCl_2_, 10 MgCl_2_, 10 NaHCO_3_, 5 trehalose, 5 sucrose, 5 HEPES-NaOH, pH 7.2. Solutions were checked with an osmometer for correct osmolarity within 5 mOsm. After stimulation, larvae were stretched, then either fixed for 25 min in 4% paraformaldehyde in HL3.1 for immunohistochemistry or transferred to electrophysiology rig for intracellular recording.

#### Larval immunohistochemistry

Fixed larval fillets were stained in primary antibody in PBX (Phosphate buffered saline pH 7.4 with 0.1% Triton X-100) with 10% normal goat serum overnight at 4°C while gently shaking. Excess antibody was removed with 3 washes in PBX for 10 min each. Secondary antibody was applied in PBX for 2 h at room temperature and then washed 3 times for 10 min each. The fillets were mounted in SlowFade Diamond AntiFade Mountant (ThermoFisher), sealed with clear nail polish and stored at 4°C until imaging. Slides were imaged within 1-2 days of mounting to preserve fluorescent signal. Antibody dilutions are as follows: Ms anti-CaMKII (#18, CosmoBio) 1:1000; Rb anti-GFP (A-11122, ThermoFisher) 1:1000; Ms anti DLG (4F3, DHSB) 1:100; Rb anti-DLG 1:1000 (Boster Bio); Rb anti-HRP (Jackson); 1:100; Alexa Fluor 647-conjugated anti-HRP (Jackson) 1:250; Alexa Fluor 488-conjugated anti-GFP (A21311, ThermoFisher) 1:250; Goat anti-Mouse Alexa Fluor 488 (A11029, ThermoFisher) 1:250; Goat anti-Rabbit Alexa Fluor 488 (A11008, ThermoFisher) 1:250; Goat anti-Rabbit Alexa Fluor 555 (A21428, ThermoFisher) 1:250; Goat anti-Rabbit Alexa Fluor 647 (A21244, ThermoFisher) 1:250; Goat anti-Mouse Alexa Fluor 647 (A21240, ThermoFisher) 1:250. Data were taken from Ib boutons at the muscle 6/7 NMJ in abdominal segments A3-4.

NMJs were imaged on either a Leica SP5 (EGFP::CaMKII or Canton S) or Zeiss LSM880 (Tra RNAi) confocal microscopes using the 488 or 633 lasers. Z stacks were acquired using a 0.5 µM step so that the entire NMJ was captured. To quantify image intensity, the Hrp channel was used to create a mask outlining the NMJ. Axons, partial boutons and type Is boutons were excluded. The mean gray value and area of the masked area was recorded.

#### NMJ mini recording

All larval recordings were performed in low calcium HL3.1 at a pH of 7.1-7.4 (70 mM NaCl, 5mM KCl, 0.8 mM CaCl_2,_ 4 mMgCl_2_, 10 mM NaHCO_3_, 5 mM trehalose, 115 mM sucrose, and 5 mM HEPES). Recordings from female and male larvae were performed after the spaced depolarization protocol (with the condition and sex blind to the experimenter). The MultiClamp 700A with the Chart5 computer software were used for recordings. A recording electrode was used to measure mEJPs in muscles 6 (45/56) or 7 (10/56) in abdominal segments A3 and A4 for 2 minutes per muscle. Electrodes had a minimum resistance of 15mΩ and an average resistance of 26mΩ. Only muscles with a muscle resistance > 5 mV and a resting membrane potential < −50 mV were included in the analysis.

Analysis was done using a custom MATLAB script (https://github.com/Griffith-Lab/NMJ_ephys_analysis/). 1 min of each recording was analyzed for mEPSP frequency and amplitude. The minimum amplitude for a mEPSP = 0.2 mV. The average membrane potential was also calculated, and recordings with more than a 5mV change in membrane potential were excluded. Resting membrane potentials were between −50 and −75 mV and were not significantly different between conditions. All statistical analysis was performed using Prism 11. 2-way ANOVA with Tukey’s multiple comparisons tests were used with a 95% confidence interval (ɑ=0.05).

### Adult brain

#### Adult brain immunohistochemistry

Adult fly brains were dissected in cold PBS and then fixed in 4% PFA solution for 30 min at room temperature. Fixed brains were washed 3 × 30 min in 0.5% Triton-PBS (PBST) solution, blocked in 10% normal goat serum solution for overnight, and then incubated in primary antibodies. Primary antibodies were removed and samples were washed in 0.5% PBST solution for 3 × 30 min. Samples were then incubated in secondary antibody overnight, then washed 3 × 30 min in 0.5% PBST solution and mounted in the Slow fade glass mounting medium. Antibody dilutions are as follows: Ms anti-CaMKII (#18, CosmoBio) 1:10,000, Chicken anti-GFP (ab13970, abcam) 1:1000, anti-chicken Alexa Fluor 488 (A11029, ThermoFisher) 1:1000, Goat anti-Mouse Alexa Fluor 647 (A21240, ThermoFisher) 1:1000, Goat anti-Rabbit Alexa Fluor 555 (A21428, ThermoFisher) 1:1000.

Adult fly brains were imaged under a 20× objective lens with the same settings using an Andor Dragonfly confocal microscope. Auto step size and settings (laser power, gain and magnification) were kept constant while acquiring the images within each individual experiment. For EGFP::CaMKII, <u>m</u>aximum intensity <u>p</u>rojection (MIP) image was generated with Z stacks encompassing the mushroom body. Mean intensity was recorded with an ROI and the values were used in the quantification. For comparing the EGFP::CaMKII intensity between male and female mushroom body across independent experiments (Figure 3C), EGFP intensities were normalized to female mean and compared with Student’s T-test using Prism 11. For whole brain intensity quantification, MIP images were generated and mean intensity was recorded from the central brain using an ROI. CaMKII quantification from mushroom body was done from a single Z slice.

#### HCR-FISH

Reagents and probes were purchased from Molecular Instruments and RNase-free water was used for all solutions. Adult female and male brains were dissected in cold PBS as previously described. They were fixed in 4% PFA for 35 min and then washed in PBST (PBS pH 7.4 with 0.5% Triton-X) two times for 20 min each. After one wash in PBS, they were incubated in probe hybridization buffer for 30 min. Overnight incubation of samples was done with probes specific to *CaMKII* mRNA (1:100 dilution, in probe hybridization buffer). Post incubation, samples were washed in probe wash buffer 3 x 15 min and then samples were incubated in amplifier buffer for 30 min at room temperature. Overnight incubation of samples was done in hairpin amplifiers (546nm: coding region; 633nm: 3’UTR probe). Finally, samples were washed in amplifier wash buffer 3 x 15 min and then mounted in SlowFade^TM^ Glass Antifade Mountant (S36917, ThermoFisher) mounting medium. The quantification for both the probes was done form whole brain (MIP image).

#### Immunoblotting

6–8-day old female and male *w^1118^* or *traRNAi* flies (mixed males and females) were frozen at −80°C. Flies were vortexed and heads were collected on dry ice using a cold sieve. The heads were then homogenized in loading buffer (4× Bolt LDS, Invitrogen, Novex with 5% β-mercaptoethanol added) and heated for 10 min. Proteins were separated by SDS-PAGE (Bolt, Bis-Tris Protein Gels, Invitrogen) and transferred to a nitrocellulose membrane (GE Healthcare). Membrane was blocked (Blocking Buffer for Fluorescent Western Blotting, Rockland Immunochemicals) and then incubated with Anti-dCaMKII Clone 18 (1:300, CosmoBio) and anti-actin mAb C4 (1:1000, Millipore). The secondary antibody used was DyLight 680 mouse. Membrane was imaged using ChemiDoc system from Bio-Rad. Intensity of bands was measured from raw images using Image J. Intensity of the CaMKII band was normalized to that of actin in the same lane. For representation brightness of the membrane is adjusted.

#### *Ex vivo* plasticity

Brains from 4 to 10 days old female and male flies were dissected in cold HL3 medium (70 mM NaCl, 115 mM sucrose, 5 mM KCl, 20 mM MgCl_2_, 1.8 mM CaCl2, 10 mM NaHCO_3_, 5 mM trehalose, 5 mM HEPES, pH 7.3) (Stewart et al., 1994). Dissected brains were transferred to and immobilized in an imaging chamber with tungsten pins. During recordings, brains were perfused with fresh HL3 saline with a flow rate of ∼2 drops per second using the 4-pinch valve and ValveLink8.3 from Automate Scientific. Glass suction microelectrodes were pulled using the p-1000 micropipette puller (Sutter Instruments) and were used to apply low amplitude electrical stimulation to the antennal lobe (AL). For each female brain, there is a corresponding male brain that was tested using the same electrode. For pre-induction and post-induction AL stimulations, 5 trains of 20 pulses at 100 Hz (pulse width of 1ms, interpulse interval of 9ms) were applied. Each train occurred at an interval of 15 s. During paired induction 12 AL stimulation trains were applied with a 5 s intertrain interval with simultaneous perfusion of HL3 + 5 µM Dopamine. The post-induction tests were run 15 min after the start of the pre-induction protocol. The isolated pulse stimulator model 2100 and the perfusion system ValveLink8.2 were used with the pClamp11 software and Digidata1550A digitizer to run the custom paired induction protocol. Images were captured at 10 frames per second using an ORCA Flash4.0 V3sCMOscamera and 40x water immersion lens on the Olympus upright microscope BX50W1. Using the HCImage Live software, images were collected with a 512 × 512 resolution and 2 x 2 binning factor. Samples were excited at 470 nm using CoolLED pE-4000 LED with Chroma 450/50excitation filter and FF01-525/45-25 emission filter.

At every frame, ΔF/F_0_ was calculated such that F is the fluorescent value at a given frame and F_0_ is the baseline fluorescent value. Peak response to each of the 5 AL stimulations pre and post induction were calculated by subtracting the average ΔF/F_0_ during the second before simulation from the peak ΔF/F_0_ during stimulation. These 5 peak responses were then averaged to calculate the average maximum responses to compare calcium influx pre and post induction. The change between responses was calculated by subtracting the pre-response from the post-response. All statistical analyses were performed using the Prism 11 software. All tests were two-tailed with a 95% confidence interval (ɑ=0.05). Individual trials for females and males were analyzed using a paired parametric T-test. Comparisons of female and male changes in ΔF/F_0_ were analyzed using a paired parametric T-test.

#### Aversive learning

Experiment was performed in an environmental room in red light at 25°C with 65% humidity (around ZT 2-5). 4-to-14-day old female and male flies were used in the assay. Flies were given at least 10 min acclimation period in room before training or testing. The learning assays were performed as described by the Quinn laboratory (Tully and Quinn, 1985). The unconditioned stimulus (US) was provided as 12 × 1 s 90 or 60 V shocks during the 1 min pairing of the US with the conditioned stimulus (CS); 10% 4-methylcyclohexanol and 3-octanol OCT were used as the odors. Flies were then given a 2 min rest. Testing for short-term memory (STM) involved 2 min of simultaneous exposure to odors, after which flies choosing either odor were counted. A performance index was calculated (number of flies choosing the conditioned odor) – (number of flies choosing the control odor)/(total number of flies). To calculate the learning index, performance indices for reciprocal trials where the identity of the control and conditioned odors were swapped were averaged.

#### Appetitive learning

Experiments were performed in an environmental room at 25°C with 65% humidity, under light phase (around ZT 2-5). Female and male flies were starved on 1% agar for 22 h a day before the training. Starved flies were given at least 10 min acclimation period in room before training. The US was provided as 2 M sucrose solution on a piece of Whatman filter paper during the 1 min 30 s CS-US pairing for STM and 2 min for long-term memory (LTM); 10% 4-methylcyclohexanol and 3-octanol OCT were used as the CS odors. Flies were then given a 2 min rest. For STM, testing involved 1 min 30 sec of simultaneous exposure to CS odors, after which flies choosing either odor were counted. A performance index was calculated (number of flies choosing the conditioned odor) – (number of flies choosing the control odor)/(total number of flies). To calculate the learning index, performance indices for reciprocal trials where the identity of the control and conditioned odors were swapped were averaged. To assess LTM, flies were transferred back in their respective agar vials and post training they were kept on standard food for 3 h. After this period, flies were transferred back to the agar vials and were maintained their till next day when they were tested and learning index was calculated. For analysis of CaMKII, female and male flies were dissected at 15 min, 1 h, 3 h and 6 h post-training. Dissected brains were stained and EGFP::CaMKII protein levels were assessed. For all timepoints, levels were normalized to EGFP::CaMKII levels recorded in untrained flies.

#### Experimental design and statistical analysis

Each assay or experiment included all possible experimental genotypes with age-matched individuals. All statistical analysis was done using GraphPad Prism 11. Presynaptic EGFP::CaMKII with HRP or postsynaptic EGFP::CaMKII with DLG recording from larval NMJs were compared using 2-Way ANOVA followed with Tukey’s Honest Significant Difference test. Membrane potential, mEJPs frequency and amplitude recording of muscle 6/7 from female and male larval fillets were also compared with 2-Way ANOVA followed with Tukey’s Honest Significant Difference test. Comparisons of CaMKII protein intensity (westerns and immunocytochemistry), EGFP intensity (for EGFP::CaMKII protein) and HCR-FISH probe intensity between female and male were done using Student’s t-test. Linear regression analysis was done on the mean EGFP::CaMKII values across timepoints (15 min, 1h, 3h and 6h) from three independent replicate experiments using GraphPad Prism 11. Slopes for male and female flies were compared using F-test. GCaMP recordings from both female and male were compared using Wilcoxon matched-pairs signed rank test. Comparison of performance index between female and male flies for short-term aversive and appetitive or long-term appetitive memory was done using Student’s T-test. Statistics and *p* value are mentioned in the legends for each figure.

## Results

### Activity-dependent increases in synaptic CaMKII are sexually dimorphic at the *Drosophila* larval NMJ

The *Drosophila* third instar larval neuromuscular junction (NMJ) has proved to be an invaluable model for the study of synaptic development, function and plasticity (He and Dickman, 2025). The excitatory glutamatergic synapses of the larval NMJ exhibit dynamic functionality using many of the same biochemical and genetic pathways that regulate plasticity in excitatory neurons in mammalian brains. Sexually dimorphic differences related to synapse area and vesicular release have been reported at this synapse (Lnenicka et al., 2006; Nijhof et al., 2016). However, whether these differences also extend to regulation of synaptic molecules like CaMKII was unknown. To address this question, we first asked if muscle 6/7 demonstrates the same type of morphological sex differences that had been reported for muscle 4 (Nijhof et al., 2016). Fig. 1A shows that male muscle 6/7 NMJs in our endogenously tagged EGFP::CaMKII line were significantly smaller in area than female NMJs.

**Figure 1.**
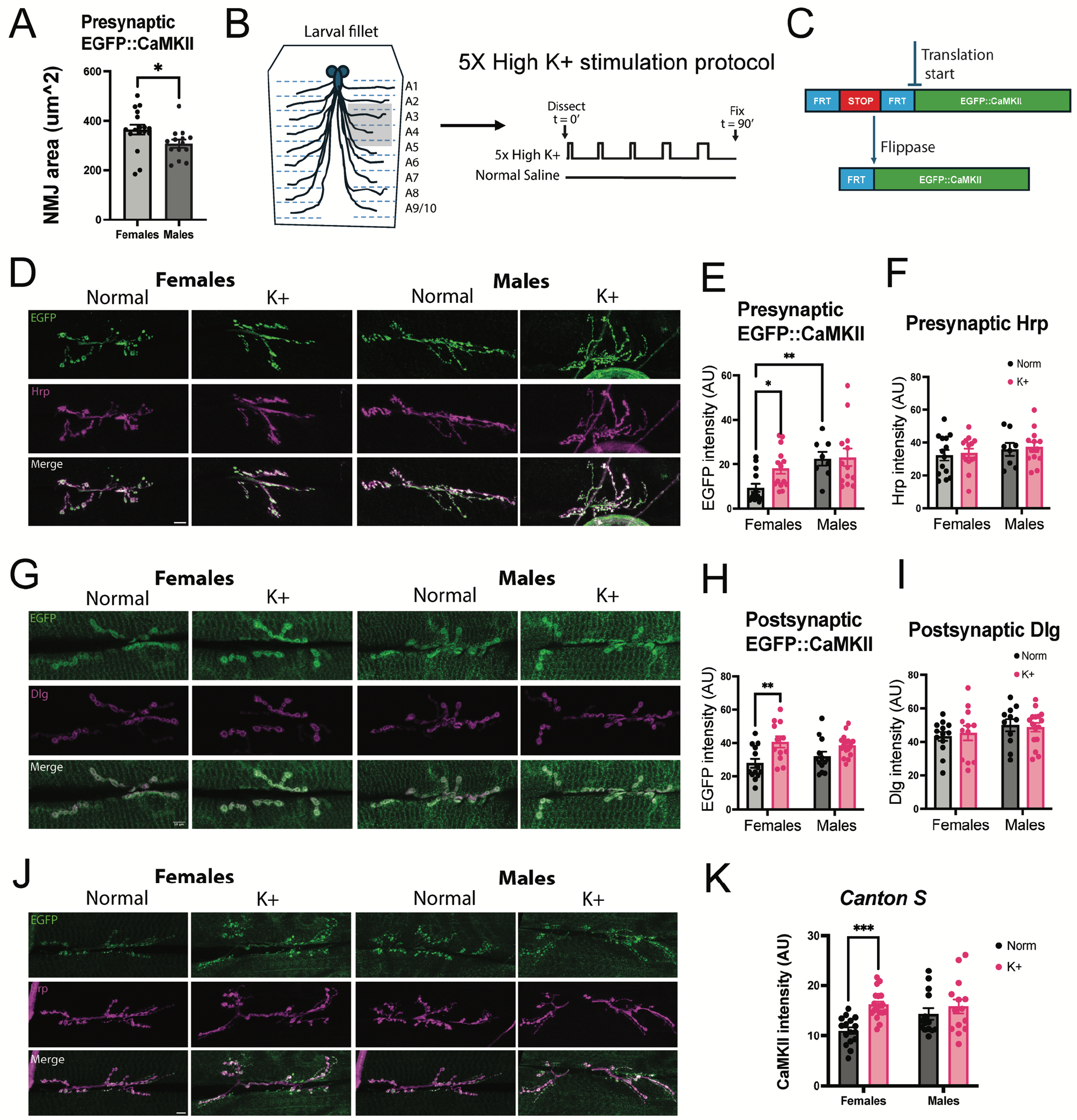
Spaced depolarization increases CaMKII protein both pre-and post-synaptically in female but not male NMJs. **A)** Bar graph representing NMJ area from female and male larvae expressing EGFP::CaMKII from the endogenous *CaMKII* locus. In comparison to males, significantly larger NMJs were observed in female larvae. (n=17 for females, n=14 for males, unpaired t-test, p=0.0427). **B)** Cartoon representation of a dissected 3rd instar larval fillet and schematic of the high potassium stimulation paradigm used to stimulate long-term plasticity. **C)** Arrangement of the *Frt-Stop-Frt-EGFP::CaMKII* cassette in the endogenous *CaMKII* locus. Expression of the tagged CaMKII occurs only in cells in which Flp recombinase is expressed. **D)** Representative images of female and male NMJs from larvae expressing presynaptic EGFP::CaMKII, subjected to high potassium (K^+^) or normal saline (Normal) washes. Fillets are stained with anti-GFP (green, to visualize CaMKII) and anti-HRP (magenta, to visualize presynaptic membrane) antibodies. Overlap shown in white. **E)** Bar graph depicting quantification of anti-GFP signal from presynaptic region of the NMJ. In comparisons to female, a significantly higher steady-state CaMKII level was observed in males. Females showed stronger changes in CaMKII levels after high potassium stimulation. n=14 for female control, n=14 for female K+, n=8 for male control, n=13 for male K^+^. p=0.0291, p=0.0058. **F)** No significant differences were observed in HRP intensity between female and male NMJ. n=14 for female control, n=14 for female K^+^, n=8 for male control, n = 13 for male K^+^. p>0.05. **G)** Representative images of female and male NMJs from larvae subjected to high potassium (K^+^) or normal saline (Normal) washes. Fillets were stained with anti-GFP (green, for CaMKII) or anti-DLG (magenta, for postsynaptic region) antibodies. **H)** Bar graph depicting quantification of anti-GFP signal from postsynaptic region of the NMJ. Females showed stronger changes in CaMKII levels after high potassium stimulation. (n=13 for female control, n=12 for female K^+^, n=13 for male control, n=16 for male K^+^. p=0.0083). **I)** No significant differences were observed in DLG intensity between female and male NMJ. (n=13 for female control n=12 for female K^+^, n=13 for male control, n=16 for male K^+^. p>0.05. All comparisons were done with 2-Way ANOVA and Tukey’s multiple comparisons post-hoc test. **J)** Representative images of NMJs from male and female *Canton S* larvae stained with anti-CaMKII (green) and anti-Hrp (magenta) antibodies. Left: control (normal saline washes); Right: high K^+^ stimulated animals. **K)** Bar graph depicting quantification of CaMKII protein levels from presynaptic region of female and male NMJs. Significant increase in CaMKII protein levels was observed in female NMJ after high K^+^ stimulation. (n=16 for female control, n=18 for female K^+^, n=14 for male control, n=14 for male K^+^. Comparisons were done with 2-Way ANOVA and Tukey’s multiple comparisons post-hoc test, p = 0.0002.

We next turned our attention to the effects of activity on synaptic proteins. Protein synthesis is required for long-lasting synaptic plasticity and many forms of long-term memory (LTM) (Frey et al., 1988; Tully et al., 1994; Martin et al., 1997). One of most important proteins involved in this type of plasticity, CaMKII, has been shown to be acutely and chronically regulated by neuronal activity in both mammals (Miller et al., 2002) and *Drosophila* (Nesler et al., 2016). To determine whether the levels or activity-dependent synthesis of CaMKII are sexually dimorphic, we exposed dissected female and male larvae to spaced, high K^+^ depolarization, a protocol that engages protein synthesis-dependent plasticity (Fig. 1B, Ataman et al., 2008) and which we have previously shown induces synthesis of both presynaptic and postsynaptic CaMKII (Clements et al., 2026). To differentiate presynaptic and postsynaptic CaMKII protein changes, we utilized Flp-Frt recombination to enable expression of an EGFP-tagged CaMKII from the endogenous *CaMKII* locus (Fig. 1C). In control larvae that were only exposed to normal HL3.1 saline, male NMJs had significantly more presynaptic EGFP::CaMKII in Ib boutons than females (Fig. 1D-E, gray bars). Interestingly, despite having lower resting levels of CaMKII protein than males, female NMJs exhibited a stronger activity-dependent increase in presynaptic CaMKII protein (Fig. 1D-E, pink bars). For NMJs expressing EGFP::CaMKII on the postsynaptic side, there was no sex difference in steady-state EGFP::CaMKII levels (Fig. 1G-H). But similar to activity-dependent changes in presynaptic CaMKII, only females exhibited robust activity-dependent postsynaptic synthesis; stimulated males had a small but non-significant increase in EGFP::CaMKII (Fig. 1G-H, pink bars).

Since it was possible that this dimorphism was specific to the EGFP::CaMKII expressing fly line, we also performed spaced high K+ depolarization in *Canton S* wildtype larvae, followed by staining with an anti-CaMKII antibody. We found that only females displayed activity-dependent CaMKII synthesis (Fig. 1J-K), confirming this is likely a general property of *Drosophila* NMJs. We also asked whether this difference in protein quantity extended generally to other synaptic proteins. We found no sex differences in either the anti-Hrp signal (Fig. 1F), which marks presynaptic glycosylated membrane proteins, or the anti-Dlg signal (Fig. 1I), which marks a postsynaptic scaffolding protein, confirming this sex difference in CaMKII synthesis is likely not due to a general difference in synaptic translation.

### Activity-dependent plasticity of spontaneous release at the NMJ is expressed in both males and females

CaMKII protein has been proposed to regulate functional synaptic changes in response to repetitive stimulation, such as the potentiation of mEJP frequency at the *Drosophila* NMJ (Kuklin et al., 2017), and the late phase of LTP in mammalian hippocampus (Miller et al., 2002). We reasoned that if activity-dependent CaMKII synthesis is different in female and males, then it’s possible that CaMKII-dependent functional plasticity may also be sexually dimorphic. Most early papers using the spaced depolarization paradigm did not specify the sex of the larvae, but several studies in which stimulation of bouton sprouting and/or increases in spontaneous release were reported explicitly indicate that male and female data were pooled (Alicea et al., 2017; San Martin et al., 2017; Maldonado-Diaz et al., 2021; Kim, 2022; Stavrovskaya et al., 2025), including one that demonstrated a requirement for protein synthesis (Lee et al., 2017) suggesting that this form of plasticity likely occurs in both sexes. To directly test whether the increase in spontaneous release can occur in males who show no increase in steady-state CaMKII levels, we recorded mEJPs from muscles 6/7 in female and male larvae after spaced depolarization. Fig. 2A shows representative traces from females and males before and after stimulation. Resting membrane potential was unaffected by either sex or treatment (Fig. 2B). Both female and male larvae showed enhanced mEJP frequency following the high K^+^ spaced depolarization protocol (Fig. 2C). Small changes in mEJP amplitude were also observed but reached statistical significance only in female larvae (Fig. 2D). These results suggest that despite differences in CaMKII accumulation, female and male larvae can both execute the same functional plasticity program to regulate spontaneous release in response to repeated stimulation.

**Figure 2.**
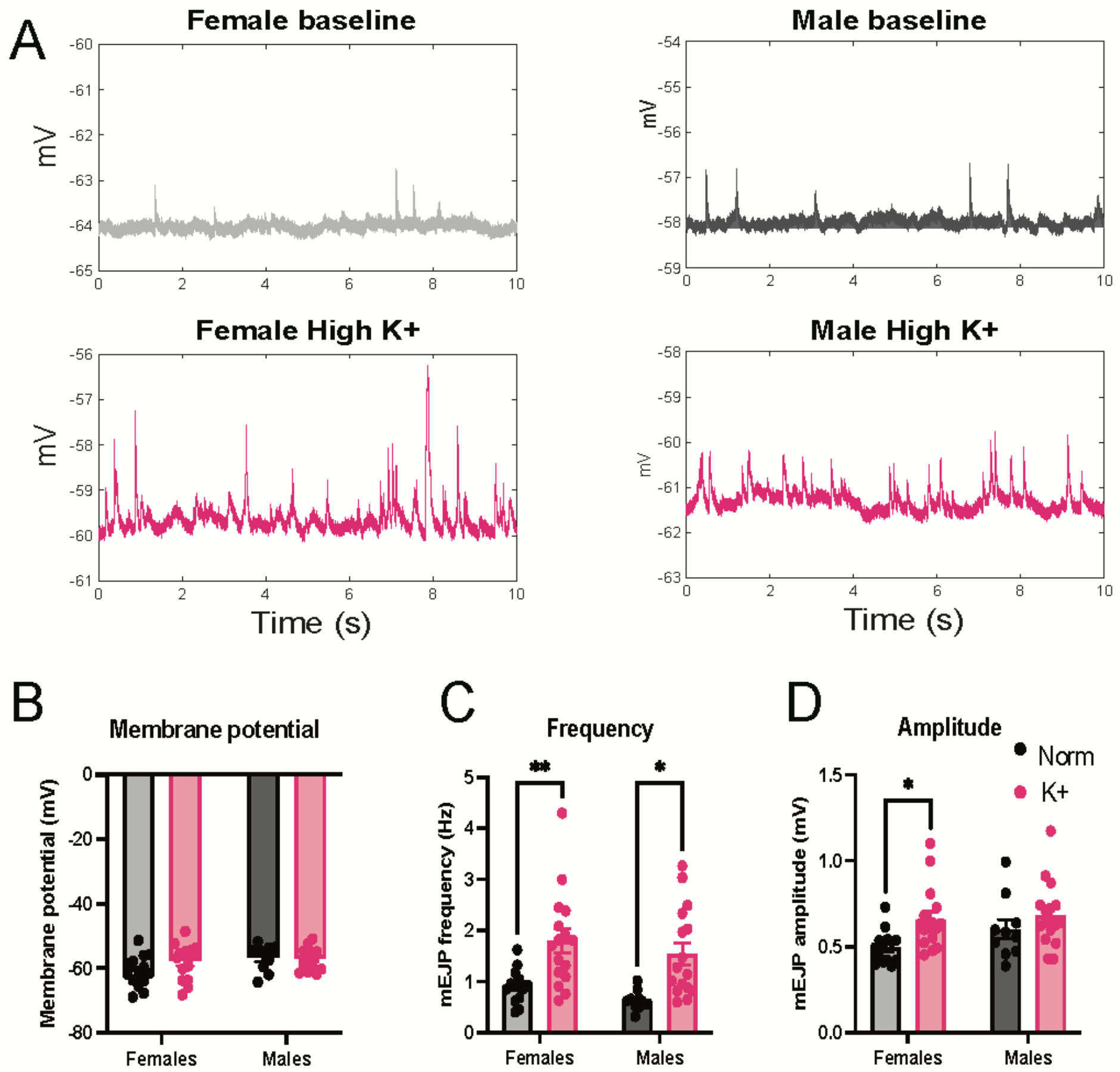
Spaced depolarization increases mEJP frequency in both females and males. **A)** Sharp electrode mEJP recordings of female and male muscle 6/7 NMJs in controls (top panels) or high K^+^ (bottom panels) conditions. **B)** Bar graph representing average resting membrane potential during recordings. Membrane potential was not significantly different between conditions or sexes. (n=14 for female control, n=16 for female K^+^, n=10 for male control, n=16 for male K^+^. For control vs. K+: females p=0.2310; males p=0.9945. For females vs. males: control p=0.0964, K+ p=0.9593. **C)** Bar graph depicting mEJP frequency. Both female and male larvae show a significant increase in rate of mEJPs after high K^+^ stimulation. Female p=0.0062; male, p=0.0109. **D)** Bar graph depicting baseline and post stimulation mEJP amplitude. Females show a significantly higher mEJP amplitude after high K^+^ stimulation. Female, p=0.0473. All comparisons were done with 2-Way ANOVA and Tukey’s multiple comparisons post-hoc test.

### Adult head CaMKII protein levels, but not mRNA levels, are sexually dimorphic

Based on our findings from NMJ, we next asked whether adult fly brain also harbors differences in CaMKII levels between female and male flies. To assess this, we first estimated CaMKII levels using immunoblotting. Homogenates of whole fly heads from *w^1118^* flies were separated by SDS-PAGE, transferred to membranes and probed with antibodies for CaMKII and actin to assess the amount of extract loaded. In line with our NMJ data, male flies had higher steady-state levels of CaMKII protein (Fig. 3A). We next asked if the observed differences in CaMKII protein might stem from a difference in mRNA abundance. To evaluate this, we assessed *CaMKII* mRNA levels using HCR-FISH. Probes for two different regions of the *CaMKII* mRNA were used, one which binds to the coding sequence of the catalytic region of the CaMKII protein (Fig. 3B left, in red) and one that binds to the distal regions of 3’UTR (Fig. 3B right, in cyan). The coding region probe is common to all *CaMKII* mRNA species, while the distal 3’UTR probe is present only on mRNAs that are synaptically translated (Chen et al., 2022). Total mRNA probe intensity was measured across the central brain. No significant difference between female and male *CaMKII* mRNA was observed with either probe (Fig. 3B, bar graph). These data suggest that the differences seen in steady-state CaMKII levels likely arise from difference in translation and/or protein degradation, rather than transcription or alterations in production of the synaptically-translated mRNA species.

**Figure 3.**
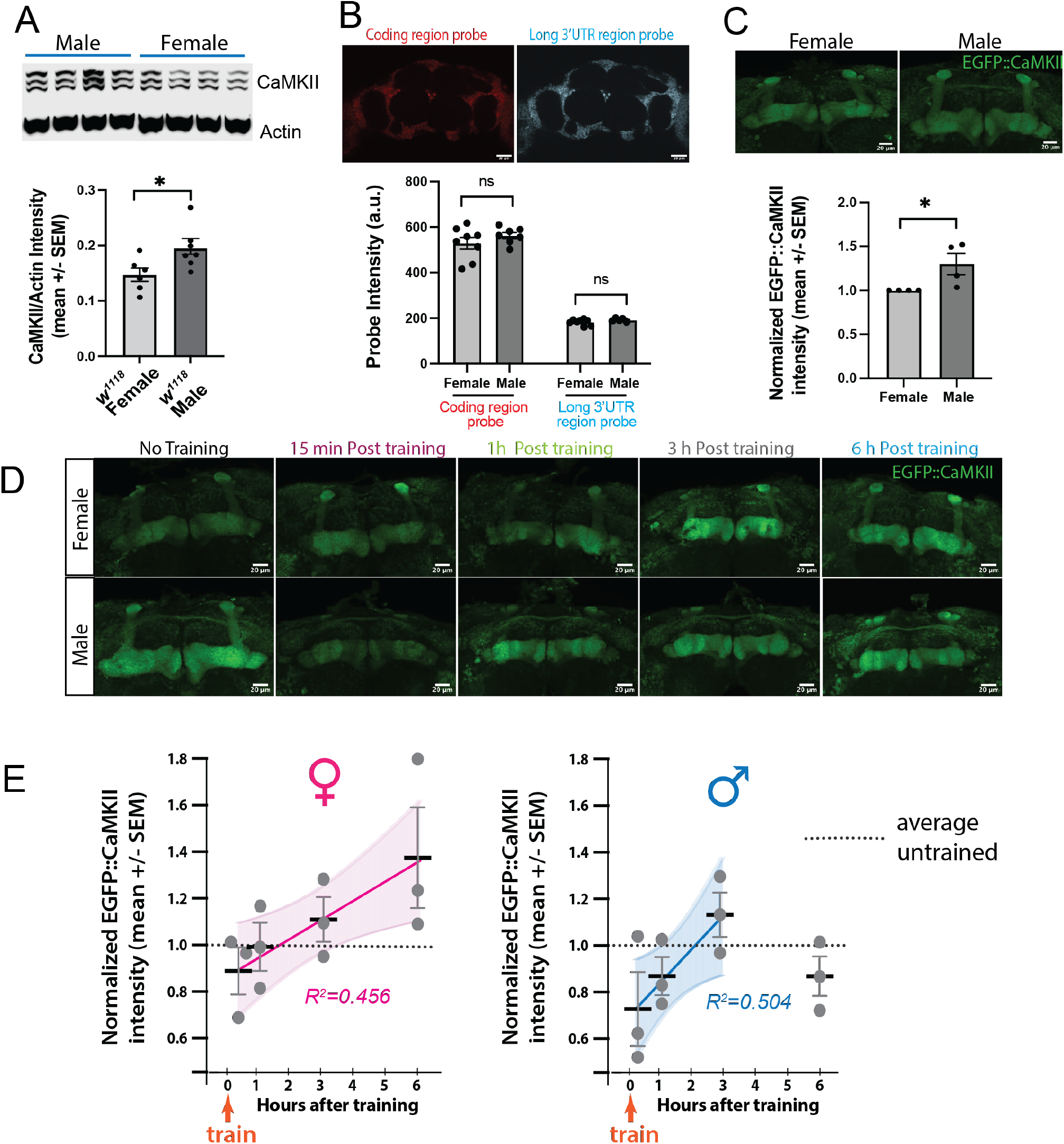
Steady-state and activity driven CaMKII levels are sexually dimorphic. **A)** Immunoblot stained for CaMKII and actin. Total CaMKII protein from female and male fly heads was quantified and normalize to actin as a loading control. Bar graph shows that female heads have significantly lower CaMKII levels than males. Comparisons done with Student T-test, p<0.05. **B)** Representative images of fly whole brain showing HCR-FISH signal for probes specific to the *CaMKII* coding region (red) and the *CaMKII* distal 3’UTR (cyan). Bar graph represents quantification of amplifier intensity for both probes No significant difference in intensity between female and male flies was observed. Comparisons done with Student T-test, p>0.05, n>6. **C)** Bar graph represents quantification of normalized GFP intensity (representing CaMKII protein) between male and female MBs from four independent replicate experiments. Mean GFP intensity value for males is normalized to female for each experiment. A significantly higher GFP intensity is observed in male flies. Comparisons done with Student T-test, p<0.05. **D)** Representative images of the MB region of female and male brains after appetitive training. Flies were dissected 15 min, 1 h, 3 h and 6 h after training. **E)** Graph representing the change in MB CaMKII protein after appetitive training. Each dot represents a complete experiment with n=5-7 brains at each timepoint. Both males and females show an initial dip in levels followed by a gradual increase. Linear regression analysis plot on mean intensity EGFP::CaMKII values from independent replicate experiments for male and female flies is shown with 95% confidence levels. Female flies show a steady increase in CaMKII protein up to 6 h post training that is not significantly different than to the first 3 h in males. Slope for female and male flies were compared using F-test, p = 0.3425. Both slopes were also significantly different from zero (F-test, female: p = 0.0159; male: p = 0.0321).

### Mushroom body CaMKII levels are regulated differently in males and females

The mushroom body (MB) is a fly brain structure housing circuits critical for learning, memory and other behaviors (Fiala and Kaun, 2024). CaMKII has long been known to be critical for plasticity in this neuropil (Griffith et al., 1993; Joiner and Griffith, 1999) and we have recently shown that the translation-dependent synaptic enrichment of this protein is required for normal synaptic plasticity and memory formation (Chen et al., 2022). To determine if female and male MBs also show baseline difference in CaMKII levels, we used our conditional expression system for EGFP::CaMKII (Fig. 1C) with a MB-GAL4 as the driver of recombination. Anti-GFP staining was quantified for MB of both female and male flies. Consistent with our head immunoblots (Fig. 3A), MBs of male flies showed higher steady-state level of EGFP::CaMKII protein (normalized to female) (Fig. 3C).

Similar to studies using high K+ at the NMJ (Nesler et al., 2016; Clements et al., 2026) CaMKII reporters in the adult brain have been shown to respond to globally applied chemical stimuli (Ashraf et al., 2006; Hillebrand et al., 2010). We were interested in determining if naturalistic stimuli, i.e. training in associative memory paradigm that produces a protein synthesis-dependent LTM, could also alter CaMKII levels and whether females and males would respond in the same way. To investigate this, female and male flies were separately trained to associate sugar with an odor. A single pairing in starved animals is sufficient to form both STM and LTM (Krashes and Waddell, 2011) with protein synthesis-dependent consolidation occurring in the hours immediately after training (Keene et al., 2006; Krashes and Waddell, 2008). To assess CaMKII protein dynamics after training, we measured EGFP::CaMKII in MB at different timepoints (15 min, 1 h, 3 h and 6 h post training) with untrained flies of the same sex as the control for normalization. Data points in Fig. 3E represent averages of individual experiments, each with n of 5-7 brains per time point.

As expected, untrained male flies started with higher levels of CaMKII protein in MB compared to females (Figure 3D, left panel). Surprisingly, at the 15 min timepoint after training, both female and male flies showed a reduction in total CaMKII protein level. The reduction was more pronounced in males than in females (Figure 3D-E, 15 min timepoint). With time, both female and male flies showed recovery in the levels of CaMKII protein, but only females had levels higher than untrained flies at 6 h (Figure 3D-E, right panel). The increase in females after 15 min was monotonic and well-fit by linear regression (Figure 3E, left panel). Regression of the complete male data set (15 min to 6 h) was poorly fit (*R^2^*=0.053, not shown). Interestingly, linear regression of the first 3 h after training in males was well-fit and had a slope that was not statistically different than that of females (F = 0.9536. DFn = 1, DFd = 17, p=0.3425). Both slopes were significantly different from zero (Female: F = 8.391. DFn = 1, DFd = 10, p =0.0159; Male: F = 7.116, DFn = 1, DFd = 7, p = 0.0321).

In broad outline, the MB data mirror what was seen at the NMJ: higher steady-state levels in males, but more robust activity-dependent accumulation in females. The difference in the initial dip in CaMKII levels suggests protein degradation may have a sexually dimorphic role in the immediate response to activity. The similar slopes over the first 3 h in males and the full-time course in females are consistent with comparable induction of protein synthesis, followed in males by earlier termination of synthesis or a second wave of degradation. However, these measurements reflect net protein accumulation; distinguishing these possibilities will require direct measurements of synthesis and degradation.

### Mushroom body-dependent plasticity is expressed in both females and males

The differences we observe in levels of CaMKII, a molecule deeply involved in synaptic plasticity in the MB, led us to compare sex-specific performance in cellular and behavioral memory formation. Previous work had established a whole-brain *ex vivo* preparation to investigate molecular and circuit-level mechanisms of memory formation (Adel et al., 2022). In this model, an explanted adult brain is exposed to artificial aversive training that mimics the standard aversive olfactory behavioral paradigm (Fig. 4A). Rather than presenting an odor as the conditioned stimulus (CS), the antennal lobe (AL) is weakly stimulated to evoke sparse activation of Kenyon cells. The unconditioned stimulus (US) is represented by a low concentration of bath dopamine since electric shock leads to release of dopamine onto the vertical lobes of the mushroom body through activation of the PPL1 cluster (Riemensperger et al., 2005; Aso et al., 2010). Paired, but not unpaired, presentation CS and US causes enhancement of GCaMP signals specifically in the α’3 lobe, but not in the α3 lobe, (Adel et al., 2022). This synaptic enhancement in α’3 replicates many features of associative learning and is believed to represent an early aversive short-term memory (STM) trace.

**Figure 4.**
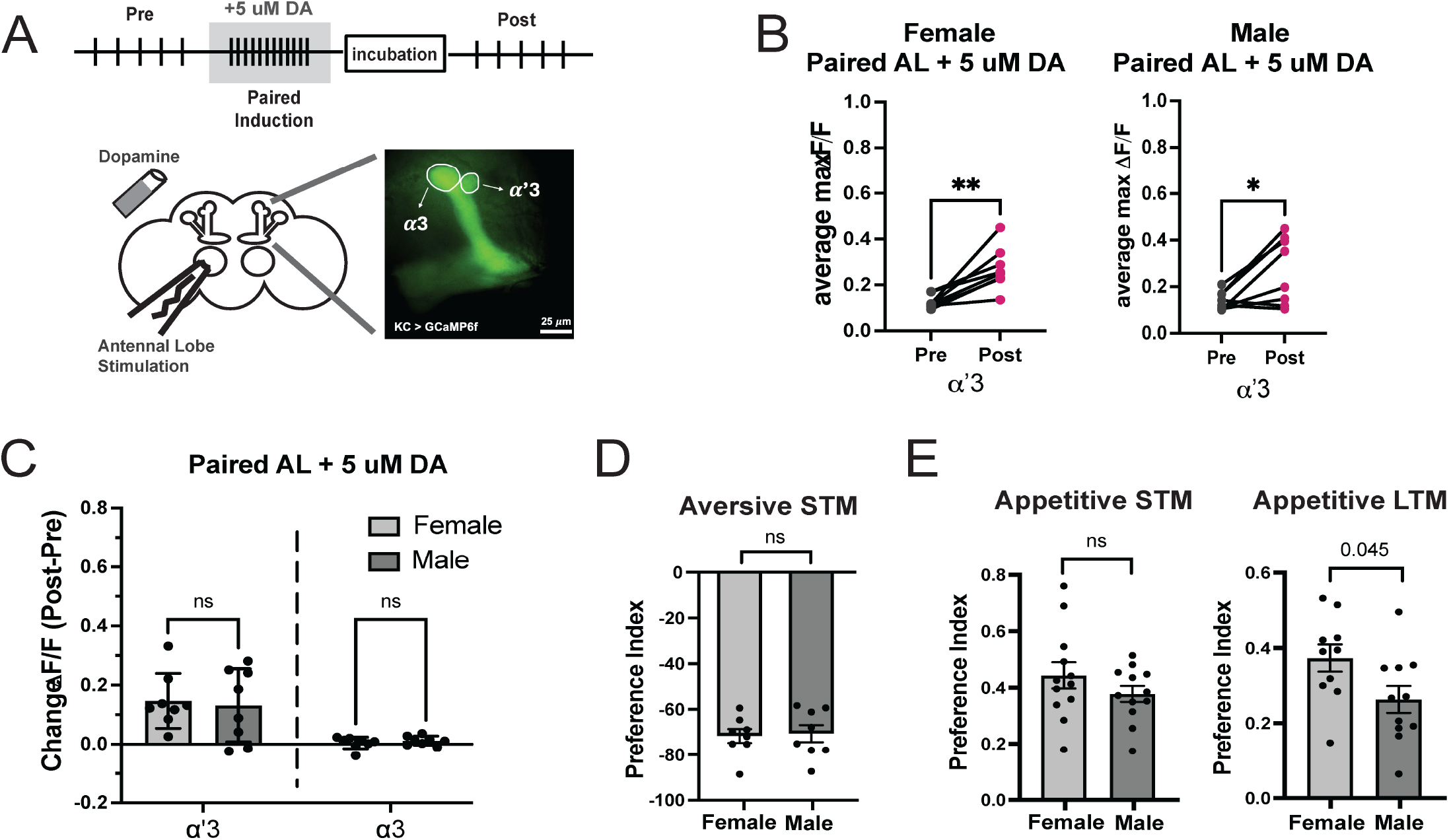
Both females and males express MB-dependent plasticity. **A)** Schematic depicting *ex vivo* paired induction protocol. **B)** Graphs depicting average max Δ*F*/*F*_0_ values from *α*’3 region of MB for both female and male flies. A significant increase in GCaMP signal was observed after induction in both female and male flies. Female, p=0.0029, n=8; male, p=0.0200, n=8. Comparisons were done with paired, parametric T-test, *α*=0.05. **C)** Bar graph depicting the change in average max Δ*F*/*F*_0_ calculated by subtracting pre-induction responses from post-induction responses. No significant difference was observed between females and males in either MB compartment. *α*′3, p=0.9797; *α*3, p=0.9973. Comparison was done with 2-way ANOVA and Tukey’s multiple comparisons post-hoc test, *α*=0.05. **D)** Bar graph depicting performance index of female and male flies for short-term aversive memory. No significant difference between female and male flies was observed. Student T-test, p>0.05. **E)** Bar graph depicting performance index of female and male flies for short-term (left) and long-term (right) appetitive memory. No significant difference in short-term memory was observed in female and male flies; however, long-term retention is stronger in female flies. Comparisons done with Student T-test, p<0.05.

Using this paradigm, we asked if there were any observable differences in cellular plasticity between adult female and male brains. We compared changes in calcium influx in response to AL stimulation before and after paired induction (Fig. 4B). Paired presentation of the US and CS causes a significant increase in calcium influx in the α’3 lobe for both females and males (Fig. 4B). The magnitude of this synaptic enhancement is not significantly different between the sexes (Fig. 4C, left). As expected, there is no significant change in the α3 lobe (Fig. 4C, right). These results indicate that associative plasticity is similarly achieved in both adult female and male brains but cannot rule out possible dimorphism in the underlying memory circuit.

To test behavioral memory directly, we trained and tested male and females separately to create odor-specific aversive (US = electric shock) or appetitive (US = sugar) memories. For the aversive paradigm we used a single training block, which produces only STM (Beck et al., 2000). In contrast, sugar training builds both STM and LTM (Krashes and Waddell, 2011). Female and male performance for both aversive (Fig. 4D) and appetitive STM (Fig. 4E, left) were indistinguishable. Interestingly, females were slightly better at forming stronger appetitive LTM for sugar (Fig. 4E, right). Whether this is due to their enhanced production of CaMKII during the memory consolidation phase or due to other factors like mating status or nutrient preference that alters circuit function (c.f. Ribeiro and Dickson, 2010; Vargas et al., 2010) remains to be determined.

### Regulation of steady-state CaMKII is downstream of the sex-determination pathway

The sex differences we have observed could arise at several levels. In mammals, circulating gonadal hormones regulate neuronal function across the brain (Gall et al., 2023). Alternatively, neurons with the same intrinsic properties might respond differently because they receive input from sex-specified upstream circuits (Nojima et al., 2010). The differences could also be cell autonomous, arising from the intrinsic sexual identity of the neurons we examine. This last possibility is particularly relevant for *Drosophila,* since biological sex determination operates primarily through a cell-autonomous genetic mechanism (Baker and Ridge, 1980). The whole-animal sexual phenotype emerges from coordination between sex-specified cells and tissues (DeFalco et al., 2008), rather than being imposed throughout the body by a circulating gonadal hormone as in mammals.

A powerful method for interrogating the role of these pathways it to short-circuit the RNA splicing cascade that establishes somatic cell sex. The ratio of X chromosomes to autosomes determines female-specific activation of the splicing factor *Sex-lethal (Sxl)*, which controls a second splicing factor, *transformer (tra)* and ultimately directs formation of sex-specific mRNAs for transcription factors that execute sex-specific gene programs. Expression of RNAi for *tra* can masculinize female fly morphology, physiology and behavior (Chan and Kravitz, 2007; Castellanos et al., 2013; Pomatto et al., 2017).

To determine whether the sex differences in steady-state CaMKII protein at the NMJ were downstream of this pathway, we expressed *tra-*RNAi using the motor neuron-specific *C380-GAL4* driver and examined anti-CaMKII levels within the area stained by the presynaptic marker Hrp. While in control NMJs, males expressed significantly more presynaptic CaMKII than females, this difference was abolished in motor neurons expressing *tra-*RNAi (Fig. 5A-B), indicating sexual dimorphism of CaMKII protein quantity is dependent on cell-specific sexual differentiation signaling originating from within the motor neuron.

**Figure 5.**
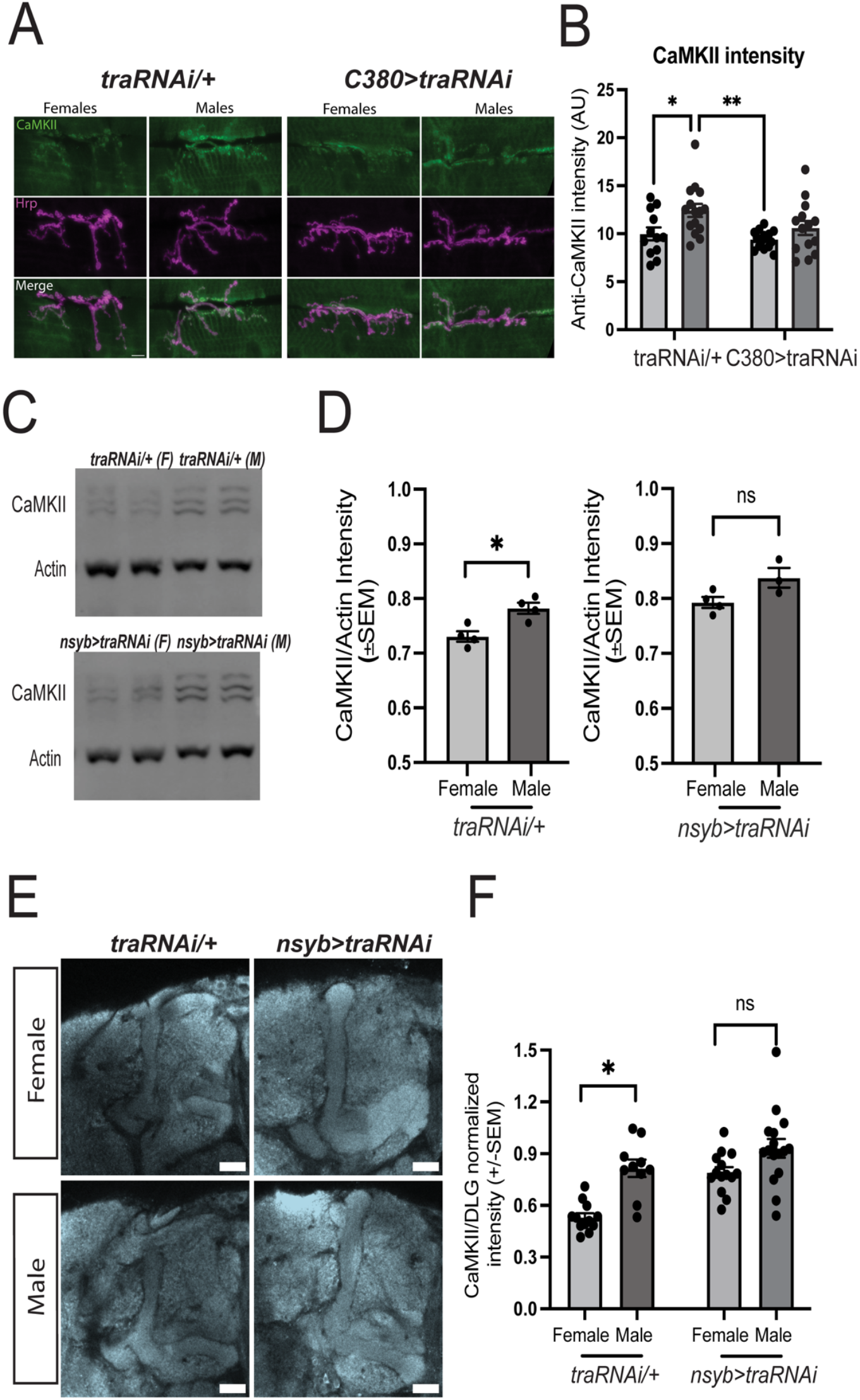
Steady-state CaMKII differences between female and male flies are controlled by *tra*. **A)** Representative images of male and female NMJs from control and *tra* knockdown larvae, stained with anti-CaMKII (green) and anti-HRP (magenta) antibodies. Overlap shown in white. **B)** Bar graph depicting quantification of presynaptic CaMKII protein. Male larvae show a significantly higher level of CaMKII protein, but the differences are abolished in *tra* knockdown lines. Control female, n=12, experimental female, n=15, p=0.036. Control male n=15, experimental male, n=14, p=0.0032. Comparisons done with 2-Way ANOVA and Tukey’s multiple comparisons test. **C)** Images of immunoblots of head samples collected from control and *tra* knockdown lines. Membrane was stained with anti-CaMKII and anti-actin antibodies. **D)** Bar graph representing intensity quantification of CaMKII protein, normalized to actin, from female and male flies for control (left) and *tra* knockdown flies (right). A significantly higher level of CaMKII protein was observed in male control flies, but the difference was abolished by *tra* knockdown. Control, p<0.05; experimental, p>0.05; n = 4. Comparisons done with Student’s T-test. **E)** Representative images (single Z section) of MB region from female and male brains for control and *tra* knockdown lines stained with anti-CaMKII antibody. **F)** Bar graph representing quantification of CaMKII protein, normalized to DLG staining, from the MB for female and male flies. A significantly higher level of CaMKII protein was observed in male MB, but the difference is abolished after *tra* knockdown. Control female, n=12; control male n=10, p<0.05. Experimental female, n=14; experimental male, n=16, p>0.05. Comparisons were done with Student’s T-test.

To examine the role of this pathway in adult brains, we downregulated *tra* gene pan-neuronally using the *nsyb-GAL4* driver. Total CaMKII protein was assessed by immunoblot from female and male fly heads for both *traRNAi/+* control and *tra* knock-down flies. In line with our previous results, a significantly higher level of CaMKII protein was observed in male control flies, but this difference was abolished when *tra* was reduced (Fig. 5C-D). We further validated these findings by quantifying CaMKII levels from MB neuropil. Similar to our observations for total protein level, MBs in male control flies showed significantly higher level of CaMKII protein then female (Fig. 5E-F). However, these differences between female and male flies were abolished with neuronal knockdown of *tra* gene. These results are consistent with the interpretation that CaMKII levels are regulated by the sexual identity of neurons.

## Discussion

Comparison of the structure of female and male brains and circuits has revealed differences in both vertebrates (Ruigrok et al., 2014; Bayless and Shah, 2016) and invertebrates (Cachero et al., 2010; Deutsch et al., 2025; Singh and Harbison, 2026). Though these neuroanatomical distinctions have been important in explaining sexually dimorphic behaviors (e.g. courtship, aggression, parental care), whether sex-specific differences also manifest at the level of the synapse and synaptic proteins remains underexplored. Using *Drosophila*, we show that male flies have a higher steady-state level of CaMKII, a protein required for induction and maintenance of synaptic plasticity, then females and that the dynamics of activity-dependent regulation of CaMKII are sex-specific. Mechanistically, these differences are downstream of *tra,* a factor in the cell autonomous sex determination hierarchy. Our findings demonstrate that sexual dimorphism extends to the molecular organization and structural plasticity mechanisms of neuronal synapses in *Drosophila*.

### A new layer of sexual dimorphism at the third instar larval NMJ

NMJs on larval body wall muscle 4 have been shown to differ in size depending on sex and genetic background (Nijhof et al., 2016). Here we show that this size difference also occurs at muscle 6/7 in our EGFP::CaMKII strain (Fig. 1A). Our immunocytochemical data add another tier of structural difference at this synapse by showing that its presynaptic terminals contain more CaMKII protein in males than in females, while anti-HRP, a marker for presynaptic membrane, is the same. Postsynaptic CaMKII and DLG, an important synaptic scaffold, were not different between sexes (Fig 1. G-I), suggesting that NMJ differences are largely due to the sex of the motor neuron.

The idea that motor neurons in *Drosophila* larvae are sexually differentiated was proposed by a study reporting motor neuron-specific differences in evoked, but not spontaneous, release on muscles 2, 4 and 5 (Lnenicka et al., 2006). Interestingly, this study did not find a difference in evoked release at muscle 6, consistent with our findings that most of the basic functional parameters at the 6/7 synapse are the same in females and males. Additionally, the ability of motor neuron-specific knock down of *tra* to masculinize levels of synaptic CaMKII in females supports the idea of cell autonomous specification of this difference. CaMKII at the *C. elegans* NMJ also appears to be specified by the sex determination hierarchy, though unlike in fly (Fig. 3B), the differential levels are achieved transcriptionally (Zeng et al., 2023).

While the structure and composition of the synapse is an important part of neuronal function, the ability of synapses to change with experience is at the heart of plasticity. To our knowledge, the effect of sex on activity-dependent changes in function has not previously been addressed at this synapse. Our study demonstrates that while both sexes can generate increases in spontaneous release in response to spaced stimulation, the molecular events underlying them differ. This mirrors work in mammalian brain showing that similar forms of plasticity can be achieved through distinct molecular pathways in the two sexes (Gall et al., 2023), and establishes the *Drosophila* larval NMJ as a genetically tractable system for dissecting the mechanisms underlying sex-dependent adaptation.

### Pathways downstream of learning are sexually dimorphic in adult *Drosophila*

There have been relatively few direct comparisons of memory performance between male and female adult flies. One study examining several forms of associative olfactory memory found no statistically significant sex differences, although females showed a trend toward better aversive LTM, similar to the modest increase in appetitive LTM we observe in females (Fig 4E; Chabaud et al., 2009). Sex differences can become more apparent under challenge. During aging, for example, males show better aversive intermediate-term memory than females, a difference associated with relative preservation of mitochondrial function (Pacifico et al., 2018). Phenotypes that emerge only under challenge support the idea that there are latent sex-dependent differences in the pathways that support plasticity. In unstressed animals, similar behavioral performance is therefore not necessarily indicative of similar underlying mechanisms.

Although sex-dependent differences in the basal activity of signaling proteins have been detected in the mushroom body (Sears and Broadie, 2022), little is known about whether core plasticity proteins are differentially regulated by learning between sexes. Our analysis of CaMKII following training directly addresses this question by examining the molecular response to induction of LTM. CaMKII levels follow different post-training trajectories in the two sexes. Thus, even when males and females show similar performance, the changes in a major plasticity protein during memory formation can be strikingly different.

### Plasticity differentially shifts the dynamics of protein accumulation in males and females

Spaced depolarization increases spontaneous release at the larval NMJ in a protein-synthesis-dependent manner (Ataman et al., 2008), and this increase is blocked in mutants that cannot generate new CaMKII due of loss of the distal *CaMKII* 3′UTR (Kuklin et al., 2017). Here, we show that both males and females express this form of plasticity, but that the accompanying regulation of CaMKII protein differs markedly between the sexes. In females, spaced stimulation produces a robust increase in CaMKII protein, whereas in males steady-state CaMKII levels remain near pre-stimulation values, raising the question of how to reconcile the need for new CaMKII synthesis with the normality of male plasticity.

Importantly, a single protein level measurement reflects a balance between synthesis and degradation, meaning that the absence of a net increase in CaMKII in males does not necessarily indicate an absence of activity-dependent CaMKII synthesis. Neuronal activity is known to stimulate local CaMKII synthesis (Nesler et al., 2016; Clements et al., 2026), but it can also activate protein degradation (Hamilton et al., 2012; Bademosi et al., 2023; Clements et al., 2026), and activated CaMKII is particularly susceptible to degradation (Quasney and Griffith, 2025). Thus, an unchanged steady-state level of CaMKII could potentially be due to increased activity-dependent protein turnover.

Our results in the adult brain support this possibility. In both sexes, training appears to produce a rapid decrease in CaMKII protein followed by recovery, consistent with an initial phase of degradation followed by increased protein synthesis. The endpoints of this program differ between males and females (Fig. 3E). In females, the initial decrease is less pronounced and the subsequent increase more robust, causing CaMKII levels to exceed baseline by 6 h after training. In males, the initial loss appears greater, while subsequent accumulation restores CaMKII only to approximately baseline. Viewed together with the NMJ results, these findings suggest that activity engages CaMKII turnover differently in the two sexes: females show net accumulation of CaMKII, whereas in males newly synthesized protein may primarily replace CaMKII lost through degradation. Similar physiological plasticity occurs despite markedly different dynamics of a core plasticity protein.

### Sex-dependent differences in steady-state CaMKII levels are downstream of *tra*

Knockdown of TRA protein in females increases resting levels of CaMKII at both larval NMJ (Fig. 5A-B) and adult MB synapses (Fig. 5E-F). Because knockdown was limited to motor neurons in larvae, this suggests that somatic sex is the determining factor for dimorphism rather than some indirect effect of sex on brain activity. In adults, pan-neuronal knockdown of TRA similarly increases CaMKII in females, but the role of brain wiring and activity is more difficult to untangle. In spite of the fact that the *CaMKII* gene has been shown to bind transcription factors that lie directly downstream of TRA in male-specific neurons (Dalton et al., 2013; Palmateer et al., 2026), transcriptional differences are unlikely to be the major mechanism causing female/male differences in CaMKII levels since steady-state mRNA levels do not differ between sexes (Fig. 3B).

A more likely set of mechanisms for differential CaMKII abundance is translation and/or degradation. Sexual dimorphism in ribosome association of mRNA has been observed in both neuronal and non-neuronal tissue in mammals (Hoye et al., 2022; Pilling et al., 2025). In *Drosophila,* TRAP from male-specific neurons immediately post-eclosion, shows that *CaMKII* is enriched in males over females by over 2-fold (Newell et al., 2016). Whether this is true in other neurons is unknown. Autophagy and proteasomal degradation have been found to be complexly regulated by stress and sex in *Drosophila* (Hansen et al., 2012; Tsakiri et al., 2013; Ratliff et al., 2016) and by sex and brain area in rodents (Du et al., 2009; Beamish and Frick, 2021; Martin et al., 2021). The particular combination of pathways underlying the differences in the dynamics of MB CaMKII remains to be determined.

In summary, we document two levels of sexual dimorphism in *Drosophila* CaMKII regulation: differences in steady-state protein abundance and differences in dynamics of protein accumulation in response to learning. Although these differences have little apparent impact on the performance of healthy animals, sexually dimorphic molecular responses can contribute to differential vulnerability to stress and disease. Even the consequences of genetic mutations can be influenced by sex-specific regulation of plasticity pathways. For example, the recent finding that a human *CAMK2A* mutation produces male-selective alterations in LTP in a mouse model (Pan et al., 2025) suggests that both basal CaMKII regulation and activity-dependent remodeling of the synaptic proteome may influence disease expression and penetrance in mammals. As the importance of biological sex in human health becomes increasingly apparent, understanding the mechanistic basis of these differences is essential, particularly given that many sexually dimorphic molecular mechanisms appear to be evolutionarily ancient and conserved (Pomatto et al., 2017; Buck et al., 2021; Regan et al., 2022; Buck et al., 2025).

## Acknowledgements

We thank Dr. Andrew Stone and the Brandeis Light Microscopy Core Facility (RRID:SCR_025892) for help with imaging.

## Author contributions

AS, KJC, ND, JB and LCG designed research; AS, KJC, ND, JB, MR and EK performed research; AS, KJC, ND, JB and EK analyzed data; AS, KJC, ND, JB and LCG wrote first draft, LCG edited paper.

